# Insights in the complex DegU, DegS, Spo0A regulation system of *Paenibacillus polymyxa* by CRISPR-Cas9-based targeted point mutations

**DOI:** 10.1101/2022.02.03.479077

**Authors:** Meliawati Meliawati, Tobias May, Jeanette Eckerlin, Daniel Heinrich, Andrea Herold, Jochen Schmid

## Abstract

Despite being unicellular organisms, bacteria undergo complex regulation mechanisms which coordinate different physiological traits. Among others, DegU, DegS, and Spo0A are the pleiotropic proteins which govern various cellular responses and behaviors. However, the functions and regulatory networks between these three proteins are rarely described in the highly interesting bacterium *Paenibacillus polymyxa*. In this study, we investigate the roles of DegU, DegS, and Spo0A by introduction of targeted point mutations facilitated by a CRISPR-Cas9-based system. In total, five different mutant strains were generated: the single mutants DegU Q218*, DegS L99F, Spo0A A257V, the double mutant DegU Q218* DegS L99F, and the triple mutant DegU Q218* DegS L99F Spo0A A257V. Characterization of the wild type and the engineered strains revealed differences in swarming behavior, genetic competence, sporulation, and viscosity formation of the culture broth. In particular, the double mutant DegU Q218* DegS L99F showed significant increase in regard to the genetic competence as well as a stable exopolysaccharides formation. Furthermore, we highlight similarities and differences of the roles of DegU, DegS, and Spo0A between *P. polymyxa* and related species. Finally, this study provides novel insights in the complex regulatory system of *P. polymyxa* DSM 365.

**Importance:** To date, only limited knowledge is available on how complex cellular behaviors are regulated in *P. polymyxa*. In this study, we investigate three regulatory proteins which play a role in governing different physiological traits. Precise targeted point mutations are introduced to their respective genes by employing a highly efficient CRISPR-Cas9-based system. Characterization of the strains revealed some similarities, but also differences, with the model bacterium *Bacillus subtilis* in regard to the regulation of cellular behaviors. Furthermore, we identified several strains which have superior performance in comparison to the wild type strain. Overall, our study provides novel insights which will be of importance in understanding how multiple cellular processes are regulated in *Paenibacillus* species.

## Introduction

To survive changing environmental conditions, complex genetic signaling networks are involved in the control of cellular adaption processes in many bacteria. In *B. subtilis*, the two-component system DegS/DegU is an important regulator for various differentiation strategies, such as motility, formation of extracellular biofilm matrixes, variation of colony architecture, synthesis of degradative enzymes, and genetic competence.^1–3^ Further adaption processes are controlled in Bacilli by the master regulator Spo0A which coordinates the transition of growing cells to spores. Besides the initiation of sporulation, this multicomponent phosphorelay system regulates transcription, directly or indirectly, of more than 500 genes involved in the adaption to nutrient starvation and changing environmental conditions.^4,5^ In combination, DegS/DegU and Spo0A are often found to either jointly or antagonistically control various adaptive traits. Morphological variations of colonies on agar plates as well as the synthesis of exopolysaccharide (EPS) or degradative enzymes, such as subtilisin, were found to be strongly dependent on the respective level of phosphorelay of both regulator systems in *B. subtilis*.^6–9^

Due to their multifunctionalities, DegU, DegS and Spo0A have been widely used as prominent targets in academia and industry for genetic optimization of Bacilli. For instance, mutations such as *degU32*(Hy) and *degS200*(Hy) were applied to stimulate synthesis of degradative enzymes with commercial significance.^1^ Moreover, deletions of the 15 C-terminal residues of Spo0A were used to block the early stage (0) of sporulation, while keeping other Spo0A-regulated genes active.^10^ Prominently, substitutions of the alanine at amino acid position 257 (A257) to either valine or glutamic acid were used to generate a sporulation-deficient strain without impairing the Spo0A-mediated *abrB* promoter repression, thus enabling increased productivity of enzymes and antibiotics as well as a delayed entry into the stationary growth phase.^11–13^

While much is known about the regulatory network controlled by DegU, DegS, and Spo0A in *B. subtilis*, only little knowledge exists about their roles in *Paenibacillus*, even though this genus has gained an enormous interest for diverse biotechnological applications from production of enzymes, EPSs, to antimicrobials and platform chemicals.^14–18^ This is, inter alia, due to the complex genetic accessibility of *Paenibacillus*,^17,19^ which limits the number of publications so far testing targeted mutations within the DegS/DegU and Spo0A systems. Therefore, most reports are limited to untargeted mutations obtained from random mutagenesis approaches or by adaptive laboratory evolutions. As an example, Hou *et a*l.^20^ detected spontaneous mutations in Spo0A (R211H) and DegU (Q218R) at once in laboratory cultivations of the biocontrol strain *Paenibacillus polymyxa* SC2, which resulted in a more stable, partially transparent morphology that was also lacking the ability to form endospores. In another study, a transcriptomic analysis of the same strain revealed a strong upregulation of *degU* and a consecutive activation of biofilm and EPS-related genes, such as *epsB, epsE*, and *abh*, during the colonization process of *P. polymyxa* SC2 in the rhizosphere of pepper plants.^21^ This is in accordance with a comparative genomic analysis between different *P. polymyxa* strains suggesting a key-role of DegS/DegU and Spo0A in the synthesis of EPSs, which in turn enables a motion and wide distribution of the strains and their antimicrobial metabolites over the plant.^22^ With regard to motility, so far the only reported targeted knockout of *degS* in *Paenibacillus* resulted in a non-motile strain as a consequence of a blocked flagellar gene transcription.^23^

To understand the impact of the regulatory systems DegS/DegU and Spo0A on adaptive traits and growth of *Paenibacillus*, we employed a CRISPR-Cas9-based system for the introduction of targeted point mutations in the respective genes. As representative strain, we have selected the industrial relevant strain *P. polymyxa* DSM 365 which has already been utilized for the production of tailor-made microbial biopolymers and 2,3-Butanediol.^15,18^ The A257V mutation in Spo0A was chosen based on its prominent role on the *B. subtilis* physiology, in order to identify the transferability to *P. polymyxa*. From this, a weakening of the promotor binding site by reduced homo-dimer formation or a decreased contact to the cognate sigma factors is expected, as observed in *B. subtilis*.^11,13^ Based on the study by Hou *et al*.^20^ which reported the double mutants of DegU and Spo0A, we have been inspired to integrate a DegU Q218* mutation in *P. polymyxa*, which mediates a premature truncation of DegU and thus potentially impairing its phosphorelay without fully abolishing its activity. In addition, DegS was modified in the DNA binding domain on position L99F, based on a report of decreased viscosity detected in a randomly mutagenized *Paenibacillus* strain having mutations in the DNA binding of DegS.^24^ Furthermore, to analyze the interaction of both regulatory systems and the impact of the mutations described, a double mutant DegU Q218* DegS L99F and a triple mutant DegU Q218* DegS L99F Spo0A A257V were generated. All engineered strains were evaluated in regard to their physiological traits as well as the changes in viscosity profile of the cultivation broth.

## Results

### CRISPR-Cas9-based targeted point mutations

In this study, the regulatory genes *degU, degS*, and *spo0A* are chosen to evaluate their roles in regulating different physiological traits and EPSs formation in *P. polymyxa* DSM 365. For *degU*, a C → T substitution at nucleotide position 652 was introduced, which led to a DegU Q218* mutant. For *degS*, a C → T substitution was targeted at nucleotide position 295 to generate DegS L99F. Meanwhile, the targeted mutation for the *spo0A* was a C → T substitution at nucleotide position 770, which resulted in Spo0A A257V. Moreover, we combined the mutations to generate the double mutant DegU Q218* DegS L99F and the triple mutant DegU Q218* DegS L99F Spo0A A257V. To achieve this, the CRISPR-Cas9-based system was employed to mediate the introduction of the desired point mutations (Figure 1A). The 20-nt spacer sequence was selected based on the closest proximity to the targeted position. For DegU, the choice of a spacer sequence in which the targeted nucleotide was located was possible. However, this was not possible for DegS and Spo0A. Therefore, in addition to the originally targeted mutations, several silent mutations were introduced in the original positions of the spacer or the protospacer adjacent motive (PAM) site in order to avoid that Cas9 attacks the desired mutants (Figure 1B). The editing efficiency varied between the different modifications. The editing efficiency for the single mutants DegU Q218*, DegS L99F, and Spo0A A257V were 4/10 (40 %), 1/20 (5 %), and 9/10 (90 %), respectively. Meanwhile, the double mutant DegU Q218* DegS L99F and triple mutant DegU Q218* DegS L99F Spo0A A257V were obtained with 1/10 (10 %) and 2/2 (100 %) editing efficiency, respectively. In this case, as observed for Spo0A A257V, it seems that a point mutation in the PAM site in combination with additional mutations in the spacer sequence increased the overall editing efficiency. Finally, the employed system was able to introduce up to five mutations simultaneously and thus would be highly applicable for generation of mutant strains with multiple targeted modifications.

**Figure 1.**
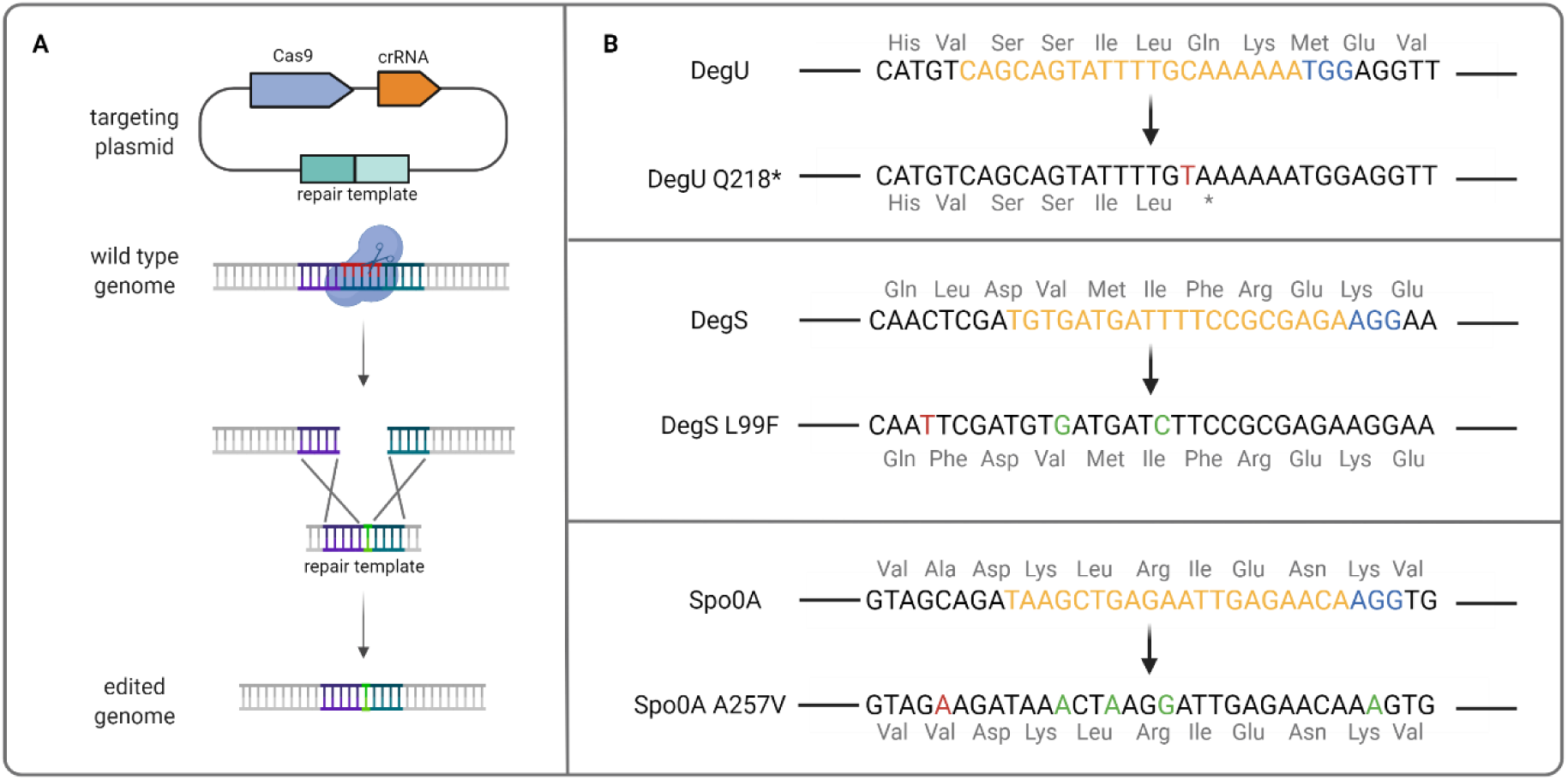
(A) Schematic overview of CRISPR-Cas9-mediated targeted point mutations. (B) Wild type sequences of *degU, degS*, and *spo0A* genes of *P. polymyxa* (top) and the respective targeted mutations in this study (bottom). Mutations which caused DegU Q218*, DegS L99F, and Spo0A A257V are indicated in red. Additional silent mutations which were added to increase the editing efficiency of the targeted point mutations are indicated in green. Spacer sequence and the respective PAM site are indicated in yellow and blue, respectively. The protein sequences of each protein are also indicated.

### Sequence alignment and protein modeling

Little is known about the role of regulatory proteins in *P. polymyxa*. Therefore, we are intrigued to see whether the proteins investigated in this study portrayed sequence similarity with the model bacterium *B. subtilis*, and if we can draw a functional correlation based on the sequence similarity, even though, for instance, *Paenibacillus* is lacking *degQ* and *degR* which are required for efficient phosphotransfer from DegS∼P to DegU and stable phosphorelay of DegU in *B. subtilis*.^23,28,29^ In general, both bacteria portrayed amino acids sequence similarities in their DegU, DegS, and Spo0A with identities of 54 %, 42 %, and 69 %, respectively. Interestingly, the targeted amino acids of DegU and Spo0A are among the conserved amino acids, which might hint the essentiality of these residues. Meanwhile, targeted L99 residue in DegS is not among the conserved amino acids but corresponds to its conservative counterparts, V102, in *B. subtilis* instead (Figure 2A, S2). In *B. subtilis*, DegU has two protein domains: the response regulatory domain (aa 5 – 121) and DNA binding domain (aa 159 – 224).^30^ The DNA binding domain of DegU is composed of helix-turn-helix (HTH) which belong to LuxR-type DNA binding domain. It is important to note that the DegU mutation in this study occurred in the DNA binding domain which leads to a premature stop codon. As mutation of DegS does not take place within the histidine kinase domain,^31^ it is not expected that the mutation will lead to significant change in its phosphorylation role. Meanwhile, Spo0A mutation occurred in the C-terminal region, away from the response regulatory domain and the HTH binding domain.

**Figure 2.**
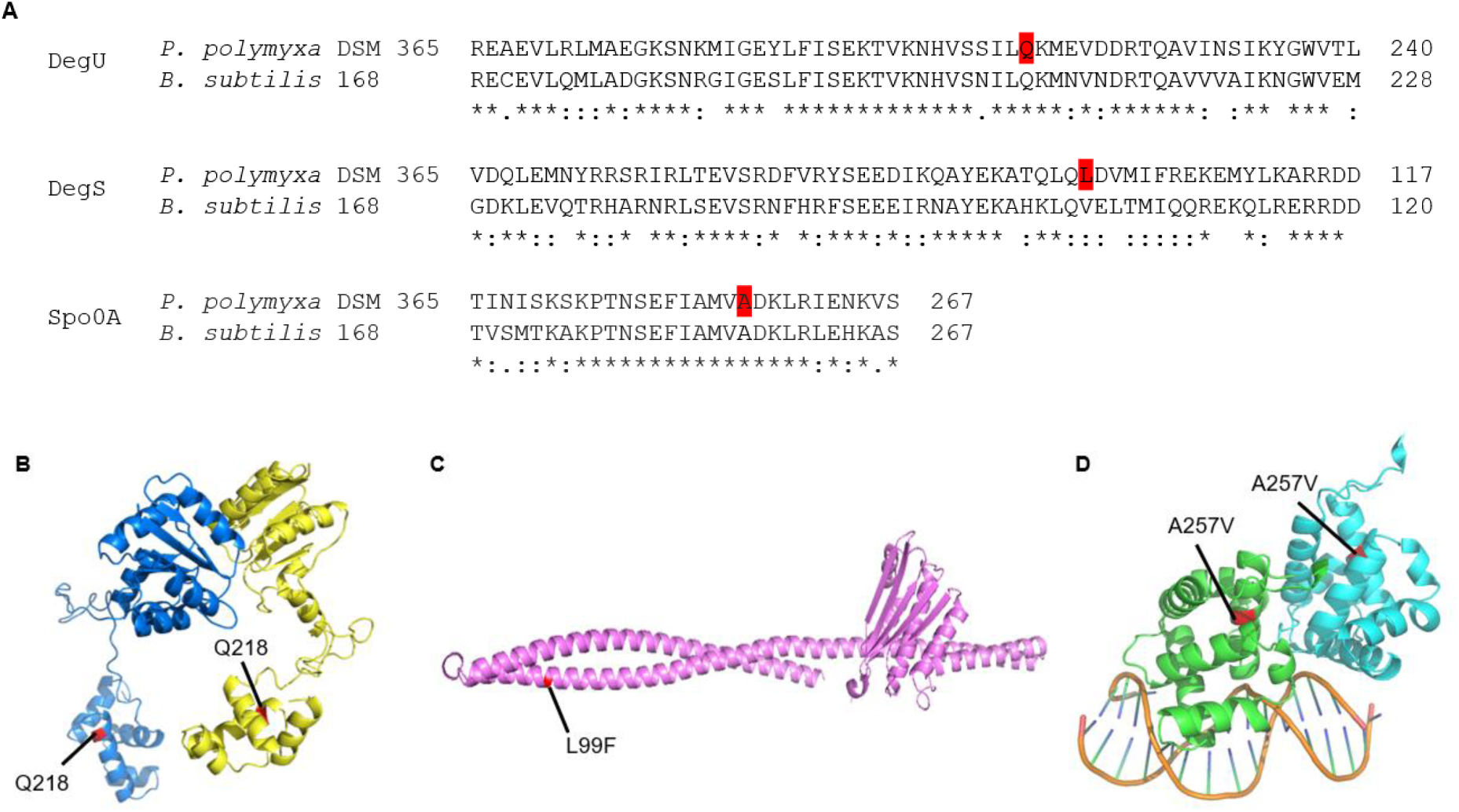
(A) Alignment of protein sequences of DegU, DegS, and Spo0A of *P. polymyxa* DSM 365 and *B. subtilis* 168. The targeted residues for the mutations investigated in this study are highlighted in red. Protein modeling of DegU (B), DegS L99F (C), and Spo0A A257V (D) of *P. polymyxa* DSM 365. DegU and Spo0A are shown in their dimer forms. For simplicity, only residue 141-267 are shown for Spo0A. The mutated residues are highlighted in red in the protein structure.

To better understand the effect of the targeted mutations to the three-dimensional structure of the proteins, we performed modeling studies for the DegU, DegS, and Spo0A (Figure 2B-2D). The Q218* mutation results in a truncated version of DegU which makes it loses one α-helix motif at the C-terminus, which might be responsible as the recognition helix. Therefore, it is plausible that this mutation would affect the binding affinity of DegU to the promoter region of the different genes it modulates. On the other hand, the L99F mutation in the DegS occurs in the coiled coil structure. As leucine is a strong helix-forming residue,^32^ its substitution to phenylalanine might weaken the helix structure in the region. Moreover, this mutation may cause a change in the phosphorylation level of the DegS itself, as observed from mutations in nearby location (S76A or S76D) in *B. subtilis*.^31^ Similarly, A257V mutation in the Spo0A is proposed to affect the flexibility and orientation of the helix structure, thus weakening Spo0A dimerization (Figure S3).^33^

### Swarming assay

In their natural habitat, it is beneficial for bacteria to have the capability to move toward nutrients or away from harmful compounds, what can ensure their survival. Likewise, *P. polymyxa* is a flagellar-forming bacterium which is capable of two types of motility: swimming in liquid and swarming on a surface.^34^ Among the engineered strains in this study, only DegS L99F retained the swarming motility while the other strains harboring mutations in DegU or Spo0A lost their swarming ability. Interestingly, the DegS L99F mutant seems to have higher swarming motility in comparison to the wild type (Figure 3). This provides insight on the importance of DegU and Spo0A in modulating swarming behavior of *P. polymyxa*. This finding is in line with what was observed for *B. subtilis*, in which knock out experiments showed that DegU is essential for swarming motility, while DegS is not.^35^ Furthermore, the previous study by Barreto *et al*.^30^ reported that a mutation at nearby position (DegU I186M) also resulted in decreased swarming motility in *B. subtilis*.

**Figure 3.**
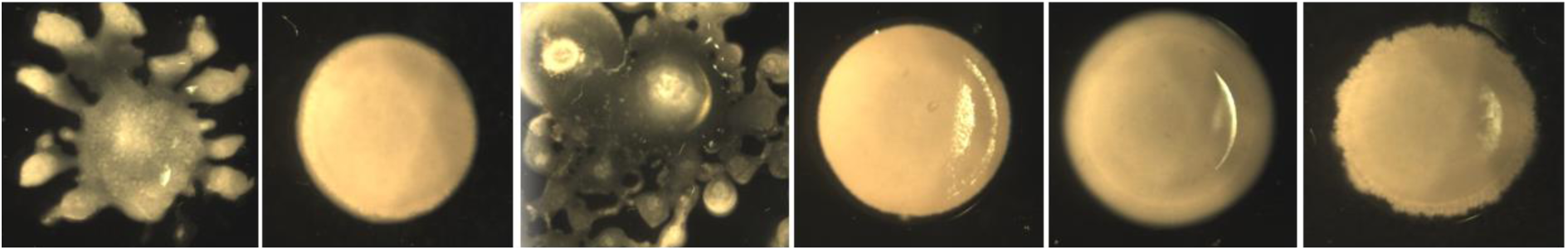
Evaluation of the swarming motility of wild type and engineered mutant strains on LB plates containing 0.4 % agar. From left to right: wild type, DegU Q218*, DegS L99F, DegU Q218* DegS L99F, Spo0A A257V, DegU Q218* DegS L99F Spo0A A257V.

### Genetic competence

Genetic competence is one of the complex cellular functions which is influenced by different cellular and environmental conditions. In this study, we investigated the role of DegU, DegS, and Spo0A on genetic competence development of *P. polymyxa*. For this, we performed conjugation between *P. polymyxa* strains and *E. coli* S17-1 harboring the non-targeting pCasPP plasmid without the HDR template. Among the five engineered mutant strains, only DegU Q218* resulted in reduced genetic competence, by almost 70 % in comparison to the wild type. Meanwhile, DegS L99F and the triple mutant DegU Q218* DegS L99F Spo0A A257V had slightly increased competence. Remarkably, the double mutant DegU Q218* DegS L99F and single mutant Spo0A A257V possessed significantly improved competence, by 19 and 13-fold higher than the wild type, respectively. While the double mutant showed substantial increase in genetic competence, the combined triple mutant of DegU Q218* DegS L99F Spo0A A257V reduced the genetic competence towards wild type level (Figure 4A). This indicates that genetic competence is regulated via a different antagonistic mechanism in *P. polymyxa* involving the DegS/DegU and the Spo0A network.

**Figure 4.**
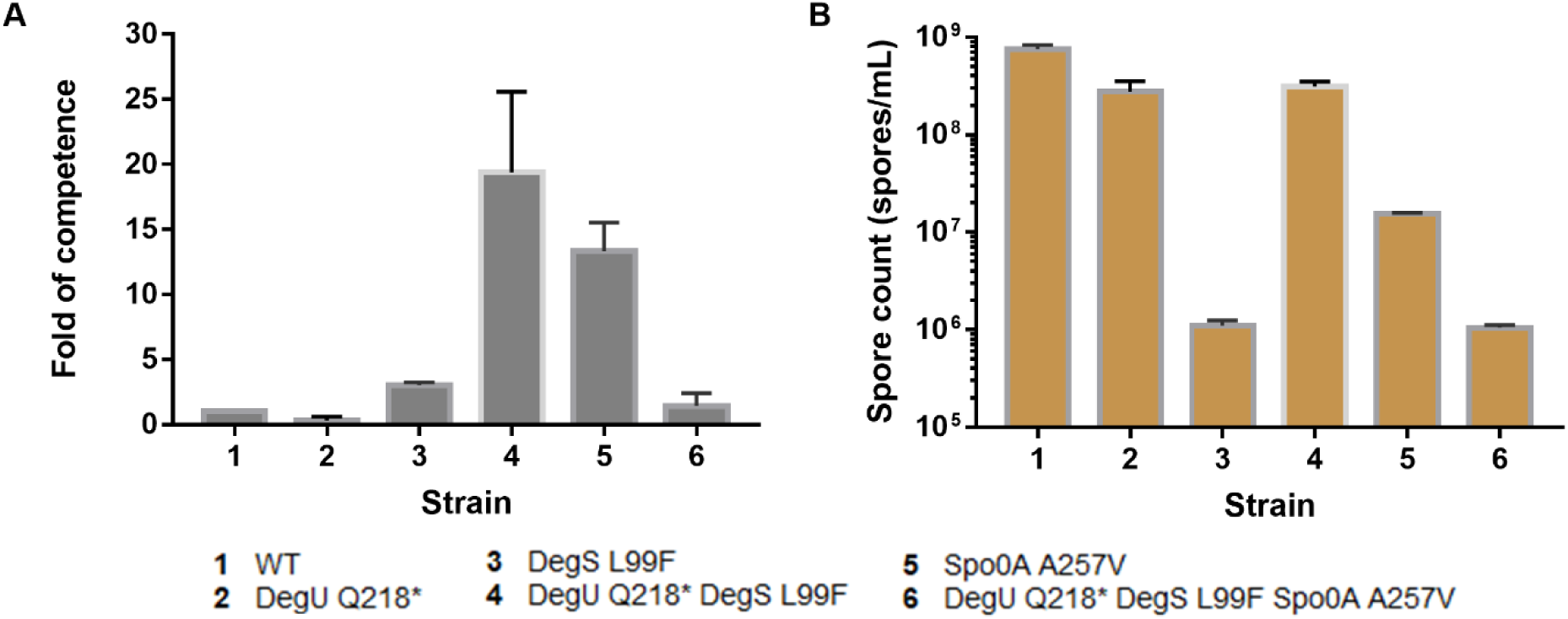
Evaluation of genetic competence (A) and sporulation (B) of the wild type and mutant strains of *P. polymyxa*. The fold of competence was calculated in relative to the wild type. Spore count of the different variants was obtained from counting the spores of the samples from the fermenters by using phase-contrast microscopy using C-Chip disposable counting chambers. The fold of competence and spore count are presented in linear and logarithmic scales, respectively. Error bars indicate the standard deviation.

In *B. subtilis*, the master regulator of genetic competence, ComK, has been studied extensively. There it was found that the non-phosphorylated form of DegU binds to *comK* and thus increases the strain’s ability for plasmid uptake.^36,37^ Furthermore, Spo0A∼P acts as an inhibitor for the competence repressor AbrB.^38^ On the other hand, no study has been reported so far on how genetic competence development in *P. polymyxa* is regulated. Genome analysis revealed that *P. polymyxa* DSM 365 carries the *abrB* gene but is lacking *comK*. From this, it can be evidenced that *spo0A* has a pivotal role in controlling genetic competence via different routes as found in *B. subtilis*, and thus might be a prominent target to optimize the genetic accessibility of *P. polymyxa*.

### Sporulation

Following the late exponential phase or unfavorable environmental conditions, some bacteria may respond the suboptimal environment by sporulation or biofilm formation. Sporulation allows the bacteria to survive in dormant state and germinate again when the condition permits. Different genes are involved in sporulation, but it is well known that Spo0A is the master regulator which regulates early stage of sporulation. Activation of the Spo0A is achieved through a phosphorylation cascade which includes a number of histidine kinases.^4^ In this study, we found that all the engineered strains can still form endospores, although the level of sporulation varies between the strains (Figure 4B). Compared to the mutants, wild type strain has the highest level of sporulation. DegU Q218* and the double mutant DegU Q218* DegS L99F have slightly lower endospore numbers in comparison to the wild type. On the other hand, the single mutant DegS L99F and the triple mutant DegU Q218* DegS L99F Spo0A A257V show 1,000-fold lower endospore formation. To our surprise, the A257V mutation of the Spo0A does not abolish the endospore formation. Primarily found in *B. subtilis*, this mutation is known to cause a weakening of the promoter binding site by reduced homo-dimer formation or a decreased contact to the cognate sigma factors^11^. Furthermore, Spo0A A257V was found to abolish sporulation via repression of either *spoIIA* or *spoIIG*, but retaining the repression of the transition state regulator *abrB* on the same level.^13^ Analysis of the crystal structure of the *B. subtilis* Spo0A found that the effect of A257V mutation was repressed by L174F and H162R. Therefore, in our case, we hypothesize that the alteration of L174 to Q174 in *P. polymyxa* in comparison to *B. subtilis* might suppress the effect of A257V mutation (Figure S2), as observed in a sporulating *B. anthracis* strain.^33^ On the other hand, the effect of DegU∼P on sporulation has been studied in *B. subtilis* in which DegU∼P influences the level of Spo0A∼P. High level of DegU∼P enhances sporulation by increasing the level of cellular Spo0A∼P.^39^ This result leads us to hypothesize that DegU mutations investigated in this study might result in lower phosphorylation of DegU, which in consequence puts the Spo0A∼P level below the threshold for sporulation initiation. This hypothesis is supported by the result of the plate-based assay, in which DegU Q218* mutation led to abolishment of the swarming motility and production of degradative enzymes (Figure 3, S4).

### Bioreactor cultivation

*P. polymyxa* is a highly promising EPSs producer bacterium which can produce both heteropolymers and homopolymer, depending on the carbon source. It produces heteropolymeric EPSs when excess glucose is available as the carbon source. On the other hand, its EPS biosynthesis shifts to the production of levan, which is a polymer of fructose, when sucrose is available as the carbon source.^14,15,40^ Production of these EPSs will result in increased viscosity of the cultivation broth. In this study, cultivation in EPS-inducing medium revealed strong differences in the growth and viscosity profiles among the different *P. polymyxa* variants. All investigated strains produced a highly viscous culture broth, but the timing of viscosity formation as well as the stability and the rheological properties were different among the strains. Remarkably, all the engineered mutants showed significantly reduced biomass formation as measured by OD_600_ compared to the wild type strain, even though the latter did use only half of the glucose provided as the carbon source (Figure 5A, 5B). Based on rheological analyses at 7/s and 100/s, DegS L99F showed the highest viscosity (Figure 5C, 5D), whereas mutants carrying a *degU* Q218* mutation showed highest viscosity at an elevated sheering rate of 1,000/s (Figure S5). Moreover, the DegS L99F single mutant, distantly followed by the wild type strain, showed the fastest viscosity formation within the first 20 h of the cultivation. However, both the wild type and DegS L99F strains initiated a strong degradation of the viscosifying matrix after 16 - 20 h cultivation time, resulting in a significant decrease of viscosity in the subsequent course of cultivation. Interestingly, when incorporating a DegU Q218* mutation into the wild type as well as in the DegS L99F mutant, the breakdown of the EPSs was mostly stopped and thus they switched toward a stable viscosity formation without further degradation. As a consequence, the final culture broth viscosity of DegU Q218* and DegU Q218* DegS L99F strains were higher compared to the wild type and the DegS L99F single mutant. On the other hand, the Spo0A A257V single mutant showed a >12 h delay in viscosity formation despite having similar growth and cultivation profile to the DegU mutant. Again, in combination with the DegU Q218* mutation, viscosity formation of the Spo0A mutant was accelerated and finally reached higher viscosity level.

**Figure 5.**
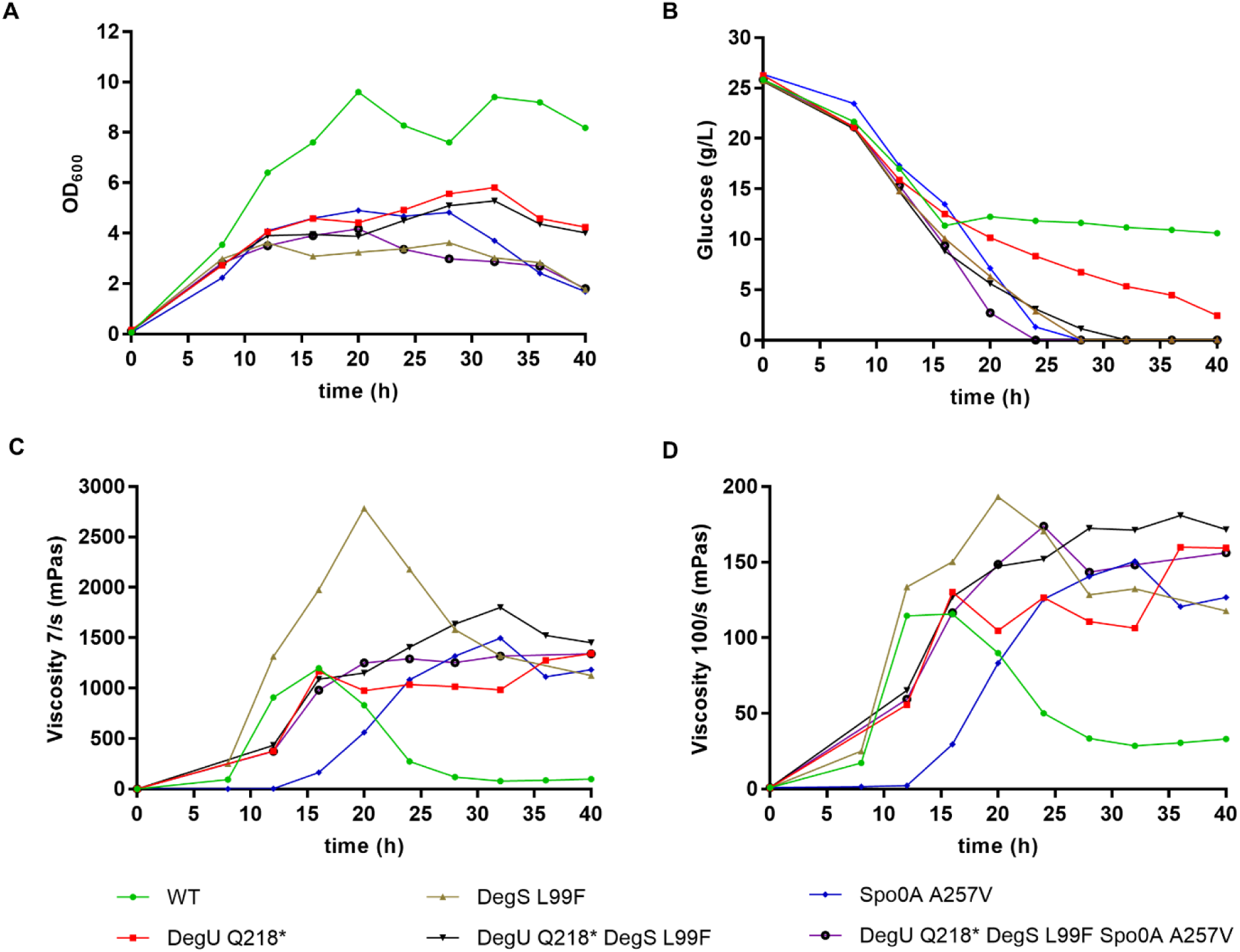
Characterization of the wild type (WT) and the engineered strains of *P. polymyxa* DSM 365 in 21 L-scale fermenters. (A) Growth profile, assessed via OD_600_ measurement. (B) Glucose consumption profile. Viscosity profile at 7/s (C) and 100/s (D) sheering rate.

## Discussion

In previous studies, we have developed efficient CRISPR-Cas-based systems to facilitate gene deletions and regulations in *P. polymyxa* DSM 365.^15,18,41,42^ Here, we demonstrated the reliability of the system to introduce targeted point mutations into the DegU, DegS, and Spo0A regulatory proteins, which are rarely investigated in *P. polymyxa*. Using the defined mutants generated by targeted integration of specific single nucleotide polymorphisms (SNPs) via CRISPR-Cas9, we propose the ability of DegU, DegS, and Spo0A in *P. polymyxa* to act as a phosphorelay-dependent multi-stage switch, which gradually controls various adaptive traits (Figure 6). In many bacteria, these three proteins are known as pleiotropic response regulators which govern multiple cellular functions. Many of these functions are under regulation of phosphorylated DegU (DegU∼P), including motility, biofilm formation, and degradative enzymes production. To date, genetic competence development is the only known cellular property which is regulated by DegU in its non-phosphorylated form.^36^

**Figure 6.**
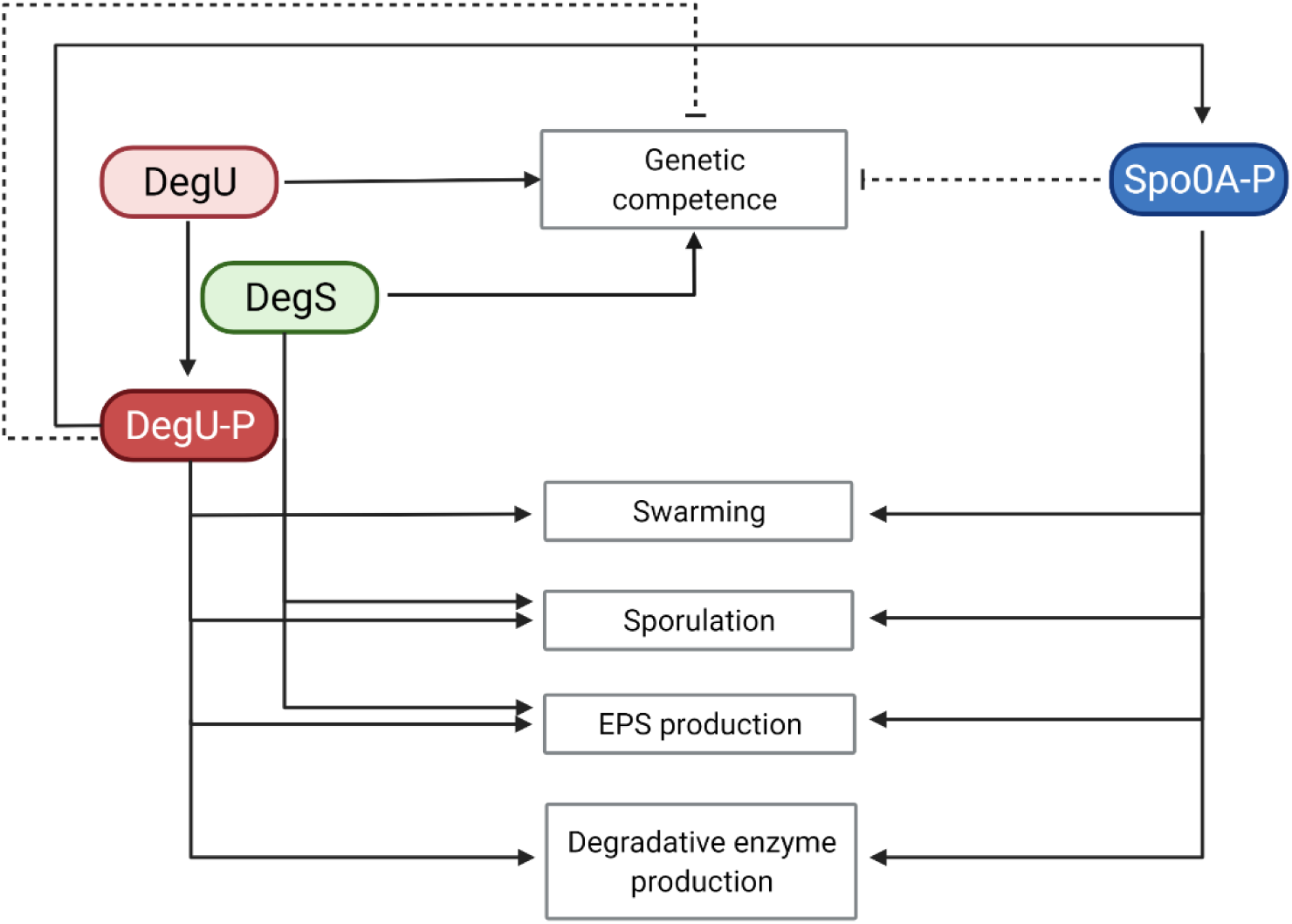
Proposed schematic model of regulatory network between DegU, DegS, and Spo0A in regulating complex cells behaviors in *P. polymyxa* DSM 365. Solid line indicates positive regulation and dashed line indicates negative regulation. Non-phosphorylated form of DegU regulates the genetic competence, while the DegU∼P is more prominent in regulating other physiological traits. Phosphorylation of DegU is catalyzed by DegS. DegU∼P positively regulates the activation of Spo0A, which in turn also regulates multiple cell behaviors.

Depending on its gradual level of phosphorylation, DegU regulates multiple cellular functions by activating their transcription via binding to the promoter regions.^2,3,43^ DegU has different binding affinities for different promoter regions, and swarming motility is among the first cellular functions that are regulated by DegU∼P.^44^ It was suggested that low level of DegU∼P is sufficient to activate flagellar biosynthesis, while hyper phosphorylation inhibits swarming motility in *B. subtilis*.^35,45,46^ In this study, it was found that Q218* mutation in DegU protein led to abolishment of swarming motility. However, this phenomenon is unlikely caused by hyper phosphorylation of DegU, considering the DegU Q218* mutant did not give positive protease assay, as it would be the case when hyper phosphorylation of DegU occurred (Figure S4).^1,43,47^ Therefore, assuming a multilevel-switch activation of genes by DegU∼P, one can speculate that DegU Q218* causes a medium level of DegU∼P because genetic competence and swarming are repressed (as expected at low levels of DegU∼P), while EPSs production is not altered (medium DegU∼P), but protease expression is not present (as expected at high DegU∼P).

In *B. subtilis*, Spo0A A257V was found to abolish sporulation, while retaining the same repression level of the transition state regulator *abrB*.^13^ As *abrB* represses the activation of the competence regulator *comK* in *Bacillus*,^38,48^ Spo0A A257V was not found to alter the genetic competence via the *abrB* route. In contrast, this particular mutation apparently did not abolish the spore formation and had a great impact on the genetic competence in *P. polymyxa*. This result particularly highlights the differences in the regulatory functions of Spo0A in *B. subtilis* and *P. polymyxa*.

Characterization of the growth and viscosity profile in the fermenters provided insights into the EPSs formation of the wild type and the mutant strains of *P. polymyxa*. To our surprise, the wild type and DegS L99F strains seem to undergo EPSs degradation, as indicated by decreased viscosity of the culture broth over the course of the cultivation process. As the EPSs from *P. polymyxa* DSM 365 are composed of glucose, galactose, fucose, mannose, and glucuronic acid as the sugar monomers,^15^ its breakdown requires the release of hydrolytic glycosidases. Transcriptomic analysis of an industrial *P. polymyxa* production strain carrying a mutation within the DNA binding domain of DegU indicated a strongly reduced expression of amylases, glucosidases, galactosidases, cellulases, levanases, as well as proteases, that may, in part, play a crucial role in the dismantling of the EPSs produced [unpublished data]. These findings are in consistence with our observations using skim milk agar plates for protease activity screening, in which the DegU and Spo0A mutant strains showed a lack of lysis zones compared to the wild type and the DegS L99F strains. Thus, a non-functional DegU regulator in *P. polymyxa* should lead to a higher, or even more stable viscosity of the broth, especially at the later stage of cultivation, as demonstrated in Figure 5B and 5C.

Unlike the DegU Q218* which resulted in decreased genetic competence, the DegS L99F strain showed 3-fold higher genetic competence compared to the wild type. From this, we conclude that the strain may have an altered phosphorelay of DegU towards lower levels of DegU∼P with the consequence of higher EPSs production without reducing the production of degradative enzymes. Moreover, in our study, the DegU Q218* DegS L99F double mutant resulted in highest viscosity formation and genetic competence, suggesting that EPSs formation requires only minor levels of DegU∼P.

Luo *et al*.^22^ has proposed two EPSs biosynthesis pathways in *P. polymyxa* based on comparative genome analysis with *B. subtilis* involving both regulatory systems. On the one hand, activation of Spo0A is mediated by autophoshorylation of *kinB* and subsequent stimulation of the *spo0F* response regulator transferring a phosphate group to Spo0A, and thus activating biofilm formation. As discussed above, it has been shown that Spo0A in *B. subtilis* negatively regulates *abrB*, which is a negative regulator of genes involved in EPSs formation.^2^ While Spo0A A257V was found to not affect repression of *abrB* expression in *B. subtilis*,^13^ it may partially repress *abrB* expression in *P. polymyxa*. As a result, EPSs formation would only occur at the later stages of cultivation, when Spo0A∼P is present at higher intracellular levels. Consequently, the Spo0A A257V mutant was found to strongly delay EPSs production during cultivation in the fermenter, while the triple mutant DegU Q218* DegS L99F Spo0A A257V showed similar early timing and levels of viscosity formation compared to the DegU Q218* DegS L99F double mutant. This indicates the presence of a second and alternative route for activation of the EPSs biosynthesis. Hence, the sensor histidine kinase DegS is triggered by an external stimulus followed by a subsequent phosphorelay of the response regulator DegU and thereby activating the biofilm genes. Since DegS/DegU promote viscosity formation through an alternative intracellular signaling pathway, their respective mutations cause strong EPSs formation that seems to cover up the altered EPSs formation through the Spo0A mutation in the same strain. From an evolutionary point of view, the regulation of EPSs formation through separate independent regulators is beneficial, as biofilm formation can have different purposes, such as nutrient concentration or the protection from harsh conditions.^49^

## Conclusion

In this study, different engineered strains were generated by introduction of targeted point mutations facilitated by CRISPR-Cas9-based system. The investigations on the pleiotropic regulators DegU, DegS, and Spo0A have provided insights on their roles in regulating different cellular response and behaviors. In particular, some of the engineered mutants are found to be superior in comparison to the wild type, particularly the double mutant DegU Q218* DegS L99F with significantly improved genetic competence. Furthermore, in contrast to the wild type, this strain does not undergo reduced viscosity of the cultivation broth which indicates stable EPSs production without any further degradation. While some similarities are observed between *P. polymyxa* and *B. subtilis*, several physiological traits seem to be regulated differently. In particular, mutation in the Spo0A which does not eliminate the sporulation and increase the genetic competence in *P. polymyxa*, which is just the opposite of *B. subtilis*. Therefore, further investigations are needed to elucidate the roles and interactions of these regulatory proteins in coordinating complex cell behaviors as well as EPS production. Finally, based on the results obtained in this study, we propose a model of the roles of DegU, DegS, and Spo0A in regulating the genetic competence, swarming motility, sporulation, EPS formation, and degradative enzymes production in *P. polymyxa* DSM 365.

## Materials and Methods

### Strains and cultivation conditions

*P. polymyxa* DSM 365 was obtained from the German Collection of Microorganisms and Cell Culture (DSMZ), Germany. Plasmid cloning and multiplication were performed in *Escherichia coli* DH5α (New England Biolabs, USA). *E. coli* S17-1 (ATCC 47055) was used as a conjugative donor strain to mediate the transformation of *P*. polymyxa. The strains were cultivated in LB media (10 g/L peptone, 5 g/L yeast extract, and 5 g/L NaCl). For plate media, additional 1.5 % of agar were used. Whenever necessary, the media were supplemented with 50 µg/ml neomycin and 20 µg/L polymyxin. *P. polymyxa* was cultivated at 30 °C while *E. coli* at 37 °C, unless stated otherwise. For liquid culture, the strains were cultivated in 3 mL of LB media in 13 mL culture tubes and incubated at 250 rpm. The strains were stored as cryo-cultures in 24 % glycerol and kept at -80 °C for longer storage.

**Table 1.** List of strains and plasmids used in this study.

| Strains or plasmids | Genotype or description | Reference |
| --- | --- | --- |
| **Strains** |  |  |
| *P. polymyxa* DSM 365 | wild type | DSMZ |
| *P. polymyxa* DegU Q218* | Q218* mutation in DegU | This study |
| *P. polymyxa* DegS L99F | L99F mutation in DegS | This study |
| *P. polymyxa* DegU Q218* DegS L99F | Q218* mutation in DegU and L99F mutation in DegS | This study |
| *P. polymyxa* Spo0A A257V | A257V mutation in Spo0A | This study |
| *P. polymyxa* DegU Q218* DegS L99F Spo0A A257V | Q218* mutation in DegU, L99F mutation in DegS, and A257V mutation in Spo0A | This study |
| *E. coli* DH5α | Cloning strain, *fhuA2* $\Delta$(*argF-lacZ*)*U169 phoA glnV44 Φ80 $\Delta$(lacZ)M15 gyrA96 recA1 relA1 endA1 thi-1 hsdR17* | NEB |
| *E. coli* DH5α::pCasPP | Cloning strain for plasmid pCasPP | 15 |
| *E. coli* DH5α::pCasPP-degU SNP | Cloning strain for plasmid pCasPP degU-SNP | This study |
| *E. coli* DH5α::pCasPP-degS SNP | Cloning strain for plasmid pCasPP degS-SNP | This study |
| *E. coli* DH5α::pCasPP-spo0A SNP | Cloning strain for plasmid pCasPP Spo0A-SNP | This study |
| *E. coli* S17-1 | Conjugation strain, *recA pro hsdR RP42Tc::Mu-Km::Tn7* integrated into the chromosome | ATCC |
| *E. coli* S17-1::pCasPP | Conjugation strain for plasmid pCasPP | 15 |
| *E. coli* S17-1::pCasPP-degU SNP | Conjugation strain for plasmid pCasPP degU-SNP | This study |
| *E. coli* S17-1::pCasPP-degS SNP | Conjugation strain for plasmid pCasPP degS-SNP | This study |
| *E. coli* S17-1::pCasPP-spo0A SNP | Conjugation strain for plasmid pCasPP Spo0A-SNP | This study |
| **Plasmids** |  |  |
| pCasPP | vector plasmid harboring SpCas9 and non-targeting crRNA, neomycin resistance (Neo<sup>r</sup>) | 15 |
| pCasPP-degU SNP | plasmid for introduction of C to T point mutation in *degU* gene at nucleotide position 652, Neo<sup>r</sup> | This study |
| pCasPP-degS SNP | plasmid for introduction of C to T point mutation in *degS* gene at nucleotide position 293, Neo<sup>r</sup> | This study |
| pCasPP-spo0A SNP | plasmid for introduction of C to T point mutation in *spo0A* gene at nucleotide position 770, Neo<sup>r</sup> | This study |

### Conjugation

Conjugation was performed between *P. polymyxa* (recipient strain) and *E. coli* S17-1 harboring the plasmid of interest (donor strain). The cryo-cultures of both strains were streaked on LB plates containing suitable antibiotics, and if necessary, following overnight liquid cultures were prepared from the colonies obtained on the plates. The overnight cultures were diluted 1:100 in 3 mL LB media, with or without antibiotic, in 13 mL plastic culture tubes. The cultures were cultivated at 37 °C, 250 rpm, for 4 h. Subsequently, 900 µl of the recipient strain was heat shocked at 42 °C for 15 min and mixed with 300 µl of the donor culture. The mixture was centrifuged at 6,000 g for 3 minutes then the supernatant was discarded. The cell pellet was resuspended in 100 µl of LB media and the resuspension was dropped on an LB agar plate. After overnight incubation at 30 °C, the cells were scrapped off the plate and resuspended in 150 µl of LB media. Afterwards, the resuspension was plated on a selection LB agar plate containing 50 µg/ml neomycin and 20 µg/ml polymyxin, according to Rütering *et al*.^15^ If necessary, the resuspension was diluted with appropriate dilution to obtain countable colonies on the plates. The plate was incubated at 30 °C for 48 h to obtain *P. polymyxa* exconjugants. Screening of the exconjugants was performed by colony PCR and sequencing of the resulting DNA fragments. Plasmid curing was performed by 1:100 subcultivation of the strains every 24 h at 37 °C.

### Plasmid construction

Targeted point mutations were realized by a CRISPR-Cas9 mediated system by use of the pCasPP plasmid as a vector base. It represents a one-plasmid system which contains the *Streptococcus pyogenes cas9*-encoding gene (SpCas9) and the respective CRISPR-RNA (crRNA) under control of the *sgsE* from *Geobacillus stearothermophilus* and *gapdh* promoter from *Eggerthella lenta*, respectively.^15,25^ To realize Cas9 targeting the different positions in *degU, degS*, and *spo0A* genes, most appropriate spacer sequences were chosen based on their closest proximity to the targeted regions. The selected spacer was located directly upstream of NGG protospacer adjacent motive (PAM) site which is recognized by the SpCas9. Approximately 1 kb homologous regions upstream and downstream of the targeted sites were amplified from the genomic DNA (gDNA) of *P. polymyxa* and provided as repair template for the homology-directed repair (HDR). Isolation of the gDNA was performed by using DNeasy Blood & Tissue Kit (Qiagen) and purification of the PCR fragments was done using the Monarch Gel Purification Kit (NEB). All fragments for plasmids cloning were amplified using Q5 DNA Polymerase (NEB) and the plasmids were assembled by isothermal assembly. Desired point mutations were introduced by the primers used for amplification of the homologous regions. For *degS* and *spo0A*, several silent mutations were also introduced in the primers to improve the editing efficiency. *E. coli* DH5α was transformed with the isothermal assembly mixture using the heat shock method. Screening of the colonies was performed by colony PCR using GoTaq Polymerase (Promega). Subsequently, the plasmids were isolated by using GeneJET Plasmid Miniprep Kit (Thermo Fisher Scientific) and sent for sequencing to confirm the correct assembly. Next, *E. coli* S17-1 was transformed with the correctly assembled plasmid, also by the use of the heat shock method. Synthesis of oligonucleotides and sequencing analysis were done by Eurofins (Germany). In silico plasmid cloning was performed by the use of SnapGene version 5.1.5. The list of oligonucleotides used in this study is provided in Table S1.

### Sequence alignment and protein modeling

Sequence alignment of the DegU, DegS, and Spo0A protein sequences of *P. polymyxa* DSM 365 and *B. subtilis* 168 was performed by using Clustal Omega.^26^ Protein modeling of *P. polymyxa* DegU, DegS, and Spo0A proteins was performed using RoseTTAFold.^27^ Visualization and analysis of the modeled proteins were done using PyMOL.

### Swarming assay

Single colonies of the wild type and mutant strains of *P. polymyxa* were inoculated into 3 mL of LB media and cultivated overnight in a shaking incubator at 30 °C and 250 rpm. The following day, 10 µl of the culture was dropped to the center of a LB plate containing 0.4 % of agar. Subsequently, the plate was sealed with parafilm and incubated at room temperature for 48 h.

### Genetic competence evaluation

Genetic competence of the different variants was evaluated by conjugating the cured strains with *E. coli* S17-1 harboring the pCasPP plasmid, following the protocol as described above. To obtain countable colonies, serial dilutions were performed before plating the conjugated strains on LB plates containing neomycin and polymyxin. The fold of competence was calculated from the total colony forming unit (CFU) of the different strains divided by the CFU of the wild type strain.

### Bioreactor cultivation

For characterization of growth properties and viscosity formation, the strains were tested in bioreactor cultivations. As preculture, the strains were cultivated in 100 mL of modified TSB medium (30 g/L TSB (Becton Dickenson), supplemented with 3 g/L yeast extract, 20.9 g/L MOPS, 10 g/L glucose) in 1 L baffled shake flasks. The preculture was grown at 33 °C and 150 rpm for 24 h. Subsequently, the precultures were transferred (1% v/v) to 21 L bioreactors (Techfors, Infors) containing 12 L MM1 P100 medium adapted from Rütering *et al*.^15^ (1.67 g/L KH_2_PO_4_, 1.33 g/L MgSO_4_ · 7H_2_O, 0.05 g/L CaCl_2_ · 2H_2_O, 5g/L peptone from soy, 30 g/L glucose monohydrate, 5 mg/L thiamine hydrochloride, 5 mg/L nicotinic acid, 0.2 mg/L riboflavin, 0.05 mg/L biotin, 1 mg/L calcium pantothenate, 5 mg/L pyridoxine hydrochloride, 0.05 mg/L cyanocobalamin, 0.05 mg/L liponic acid, 13 mg/L MnSO_4_ · H_2_O, 4 mg/L ZnCl_2_, 4.6 mg/L CuSO_4_ · 5H_2_O, 2.8 mg/L Na_2_MoO_4_ · 2H_2_O, 15 mg/L Fe_2_(SO_4_)_3_ · H_2_O, and 0.4 g/L citric acid). Cultivation was performed at 30 °C for 40 h, pH was set to 6.8 and adjusted with H_3_PO_4_ (25 %) and NaOH (1 M). In the bioreactor, target dissolved oxygen level was set at ≥ 30 % in a stirrer-gas flow cascade. To prevent sheering of the EPSs produced, agitation was limited to 300 – 600 rpm while using a stirrer setup consisting of two propellers and one Rushton, in which the latter was placed near the agitator shaft. To maintain oxygen supply, aeration was performed at 5 – 30 L/min at 0.5 bar pressure. Struktol J673 was used as antifoam agent. Samples of the culture broth were taken every 4 h. Growth, sporulation, and viscosity were assessed from the whole culture broth sample, whereas sugar consumption was analyzed from the supernatant obtained after 5 min centrifugation at 13,000 g.

### Spores quantification

Spore counts in fermentation samples were evaluated by phase-contrast microscopy using C-Chip disposable counting chambers (Neubauer/NanoEnTek) according to the manufacturer’s manual. For accurate counting, fermentation samples were serially diluted with sterile 0.9 % NaCl solution. Generation of dilution series and counting of spore titer was done in triplicates for each sampling point.

### Sugar profiling

Glucose consumption over the course of the cultivation in bioreactors was measured by HPLC equipped with an RI detector and Aminex HPX-87 H, 300 × 7.8 mm column (Bio-Rad) at 30 °C, 0.5 mL/min eluent flowrate (5 mM H_2_SO_4_), and 30 min runtime. For analysis, 2 mL fermentation samples were centrifuged for 5 min at 13,000 g, then supernatant was filtered through a 0.2 µm membrane before used for the HPLC analysis.

### Rheological analysis

Rheological analysis of culture broth viscosity was conducted every 4 h over the course of the cultivations using an Anton Paar MCR302 rheometer with double slid geometry (Measuring Cup: C-DG26.7/SS/Air, temperature: 30 °C, sample volume: 5 mL of whole culture broth). The samples were preconditioned in a pre-shear experiment at constant shear rate of 10/s for 100 s. 10 data points were recorded every 10 s. After preconditioning, viscosity was measured as a function of the shear rate. Therefore, the shear rate was logarithmically increased from 1/s to 100/s while logging a total of 25 data points.

## Acknowledgments

The authors would like to thank Christa Teckentrup for her excellent technical assistance. This study is supported by BASF SE, Germany. MM, TM, DH, AH, and JS designed the experiments and wrote the manuscript. MM and JE performed the experiments. All authors contributed in data analysis. All authors read and approved the final manuscript.

